# GeneChat: A Multi-Modal Large Language Model for Gene Function Prediction

**DOI:** 10.1101/2025.06.05.658031

**Authors:** Shashi Dhanasekar, Akash Saranathan, Pengtao Xie

## Abstract

Accurately predicting gene function from DNA sequences remains a fundamental challenge in genomics, particularly given the limited experimental annotation available for most genes. Existing computational approaches often formulate function prediction as a classification task over predefined categories, limiting their flexibility and expressiveness. We introduce GeneChat, a multi-modal large language model designed to generate free-form, natural language descriptions of gene functions directly from nucleotide sequences and textual prompts. GeneChat integrates three components: a DNABERT-2-based gene encoder optimized for long-range genomic context, an adaptor that aligns gene representations with the input space of a large language model, and Vicuna-13B, a fine-tuned LLaMA-2 variant used to produce coherent functional narratives. Trained on over 50,000 genes from the NCBI database, GeneChat outperforms GPT-4o on BLEU and METEOR metrics, demonstrating superior ability to generate accurate, context-aware, and semantically rich descriptions. This work highlights the potential of foundation models for advancing interpretable and scalable gene function prediction in a free-form, language-driven paradigm.

## Introduction

Elucidating the function of genes is a fundamental challenge in modern biology and bioinformatics. Advances in genome sequencing have yielded an explosion of new gene sequences, yet the vast majority of these genes remain uncharacterized. In fact, less than 1% of known protein sequences have any experimental functional annotation, and even in well-studied model organisms, the majority of genes have no assigned function (1). This growing gap between the number of sequenced genomes and the number of genes with known functions highlights an urgent need for improved computational gene function prediction methods (2). Robust gene function predictions can guide experimental studies, helping to prioritize genes for laboratory characterization and advancing our understanding of biological processes (2).

Traditional computational approaches to gene (and protein) function prediction have relied on a combination of sequence analysis and other biological data. Common strategies include sequence homology searches (e.g., BLAST (3), which transfers annotations from known genes to similar unknown genes), co-expression network analysis (4), protein–protein interaction networks (5), and gene ontology (GO) (6) enrichment or other classification methods (2). While these approaches have been valuable, they also face significant limitations (2). Many current methods formulate function prediction as a classification task — assigning genes to one or more predefined categories or ontology terms (1). This discrete, label-centric view fails to capture the nuanced and context-specific nature of gene functions (7). Important details can be lost when a gene’s role is compressed into a few broad categories. Moreover, classification-based models are typically restricted to predicting from a fixed set of known functions, making it difficult to describe novel functions or provide insights beyond those categories. As a result, conventional methods often produce incomplete or overly generic annotations, and their accuracy has plateaued in recent community-wide assessments (1).

Large language models (LLMs) (8) offer a promising new direction to overcome these limitations. LLMs like GPT-style transformers (9) have demonstrated an ability to encode vast amounts of knowledge and generate coherent, explanatory text in natural language. Recently, researchers have begun exploring LLMs for biological applications (10). For example, Huo et al. introduced ProteinChat, a multi-modal LLM that takes a protein’s amino acid sequence as input and generates a comprehensive narrative describing its function (7). This approach showed that free-form text generation can capture the nuanced, complex nature of protein function far better than classification, producing rich descriptions within a unified framework (7). Inspired by these advances in the protein domain, we propose to bring LLM-based, multi-modal modeling to gene sequences. By allowing an AI model to write out a gene’s functional characteristics in sentences and paragraphs, we can obtain predictions that are more flexible and informative than a list of GO terms or a binary function yes/no call.

Here we introduce GeneChat, a multi-modal large language model designed for gene function prediction from a nucleotide sequence. GeneChat accepts two inputs: (1) a DNA sequence (e.g., a gene’s coding sequence) and (2) a textual prompt (such as “Predict the function of this gene”). Given these, it produces a free-form, natural-language description of the gene’s likely function. GeneChat’s architecture consists of three key components working in tandem: (i) a gene encoder that reads the input nucleotide sequence and extracts a learned representation of that gene; (ii) an adaptor module that bridges the sequence representation into the LLM’s latent space; and (iii) a pre-trained large language model that generates the functional description conditioned on both the gene representation and the prompt.

The gene’s sequence features are injected into the language model, which then “translates” those features into a human-readable functional summary. Importantly, by using an adapter strategy, we can integrate sequence information without retraining the entire language model, leveraging the rich semantic knowledge already present in LLMs (10, 11). The result is a single unified model that can answer diverse questions about a gene’s function in natural language, much like an expert chatting about the gene.

The key novelty of GeneChat lies in its ability to produce free-form, detailed functional predictions instead of choosing from predefined categories. Unlike prior gene function predictors that output discrete labels or GO terms, GeneChat can elaborate on a gene’s role in paragraph form, mentioning possible molecular functions, biological processes, cellular locations, and even caveats or confidence levels in its predictions. This level of detail provides richer insight that can be invaluable to researchers. For instance, if a gene is predicted to be an enzyme, GeneChat might describe the reaction it catalyzes, the pathway it participates in, and analogies to well-studied proteins, rather than simply tagging it as “kinase” or “transferase.” Such comprehensive narratives make the predictions more interpretable and informative, bridging the gap between raw bioinformatics output and the kind of description one might find in a textbook or review article.

## Results

### GeneChat overview

GeneChat is a multi-modal large language model designed to enable free-form prediction of gene functions. It accepts as input the nucleotide sequence of a gene and a textual prompt (e.g., “Predict the function of this gene”), and produces a detailed natural language description of the gene’s likely function (Fig. 1a). The architecture of GeneChat comprises three main components: a gene encoder, a large language model (LLM), and an adaptor that bridges the two. The gene encoder processes the nucleotide sequence to extract a representation vector, while the adaptor ensures compatibility between the gene embedding and the LLM’s latent representation space. The adapted gene representation, together with the textual prompt, is then passed to the LLM, which generates a free-form textual prediction. For gene encoding, GeneChat employs DNABERT-2 (12), a pretrained transformer model (9) specifically optimized for genomic sequences and capable of modeling long-range dependencies within DNA. For the language generation component, GeneChat utilizes Vicuna-13B (13), a fine-tuned open-source LLM derived from LLaMA-2 (14), known for its strong performance in instruction-following and dialogue tasks.

**Fig. 1.**
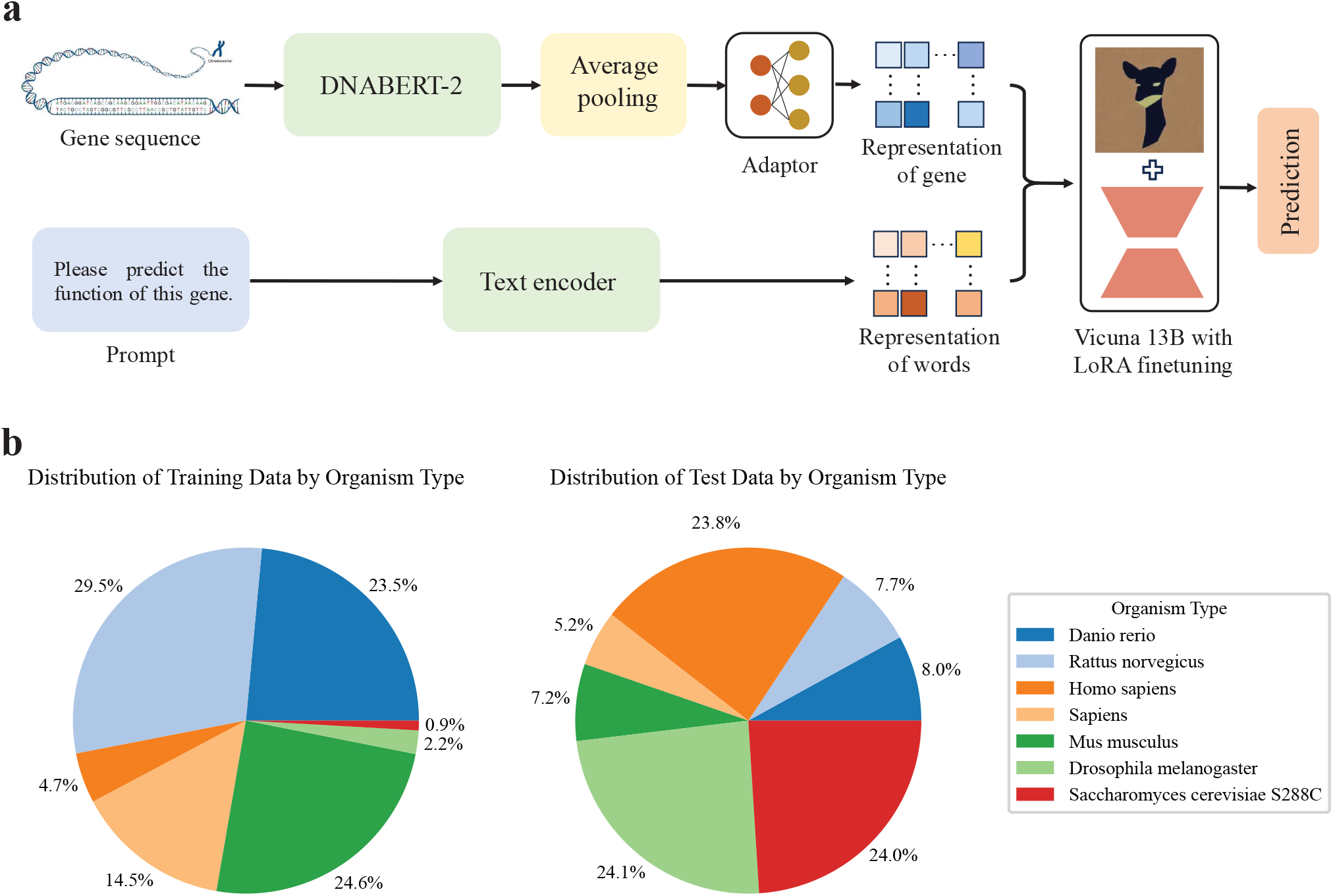
GeneChat overview. **a**, GeneChat is a multi-modal large language model developed to enable free-form prediction of gene functions. It takes as input the nucleotide sequence of a gene along with a textual prompt (e.g., “Predict the function of this gene”), and generates a detailed natural language description of the gene’s likely function. The architecture consists of three main components: a gene encoder, a large language model (LLM), and an adaptor that bridges the two. The gene encoder processes the nucleotide sequence to produce a contextual embedding, while the adaptor transforms this embedding into a representation compatible with the LLM’s input space. The adapted gene representation is then combined with the textual prompt and passed to the LLM, which generates the corresponding functional description. **b**, To train and evaluate GeneChat, we curated a dataset from the National Center for Biotechnology Information (NCBI) database, comprising 50,248 genes drawn from a wide range of organisms. Each entry includes the gene’s nucleotide sequence, a detailed textual description of its function, and additional metadata such as gene type and chromosomal location. The nucleotide sequence serves as the model input, while the function description is used as the output target during training. The dataset was randomly split into training and test sets using a 95:5 ratio.

To train and evaluate GeneChat, we curated a dataset from the National Center for Biotechnology Information (NCBI) database (15), comprising 50,248 genes from a diverse range of organisms (Fig. 1b). Each entry includes the nucleotide sequence of the gene, a detailed textual description of its function, and additional metadata such as gene type and chromosomal location. The nucleotide sequence is used as the model input, while the corresponding function description serves as the output target during training. The dataset was randomly partitioned into training and testing subsets using a 95:5 split.

### GeneChat outperforms GPT-4o in free-form gene function prediction

We evaluated GeneChat’s ability to generate free-form descriptions of gene functions by comparing its outputs on the test set against ground truth annotations using BLEU (16) and METEOR (17) scores. BLEU-1 measures unigram (single-word) overlap, while higher-order BLEU scores assess the correctness of longer n-gram sequences, capturing fluency and phrase-level accuracy. METEOR complements BLEU by incorporating both precision and recall, as well as semantic features such as stemming and synonym matching. It also penalizes disordered output via a fragmentation penalty, making it more sensitive to word order and meaning alignment. We compared GeneChat with GPT-4o, a state-of-the-art general-purpose large language model. In our evaluation, we provided GPT-4o with the name of a gene and prompted it using the following instruction: “Given the name of the gene: [*Gene Name*], predict the function of this gene.” We also tried providing GPT-4o with the full nucleotide sequence of a gene as input, it failed to generate meaningful functional predictions.

GeneChat consistently outperformed GPT-4o across all evaluation metrics (Fig. 2). It achieved a BLEU-1 score of 0.1937, substantially higher than GPT-4o’s score of 0.1444. This performance gap widened with higher-order BLEU scores: GeneChat obtained BLEU-2, BLEU-3, and BLEU-4 scores of 0.1384, 0.1065, and 0.0816, respectively, compared to GPT-4o’s 0.0563, 0.0208, and 0.0088. The pronounced difference in BLEU-4 highlights GeneChat’s enhanced ability to generate longer, more contextually accurate sequences. In addition, GeneChat achieved a METEOR score of 0.2725, outperforming GPT-4o’s 0.2422. Given that METEOR accounts for synonymy, stemming, and word order, this improvement reflects GeneChat’s superior capacity to produce responses that are not only content-aligned but also semantically coherent and fluently structured. Fig. 3 presents two examples in which GeneChat’s predictions closely align with the ground truth annotations.

**Fig. 2.**
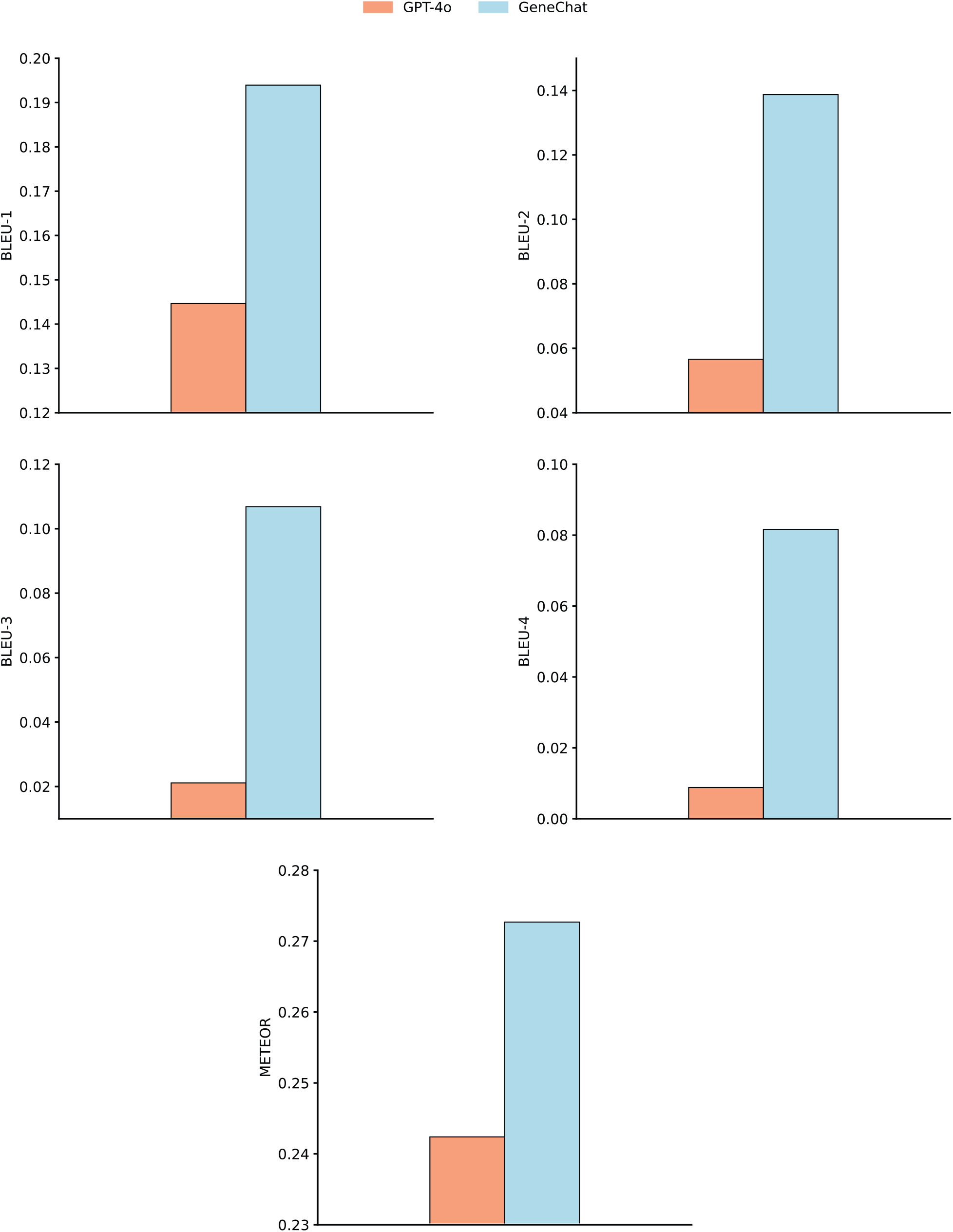
GeneChat outperforms GPT-4o in free-form gene function prediction. GeneChat significantly outperforms GPT-4o in generating free-form descriptions of gene functions, as evaluated by BLEU and METEOR scores.

**Fig. 3.**
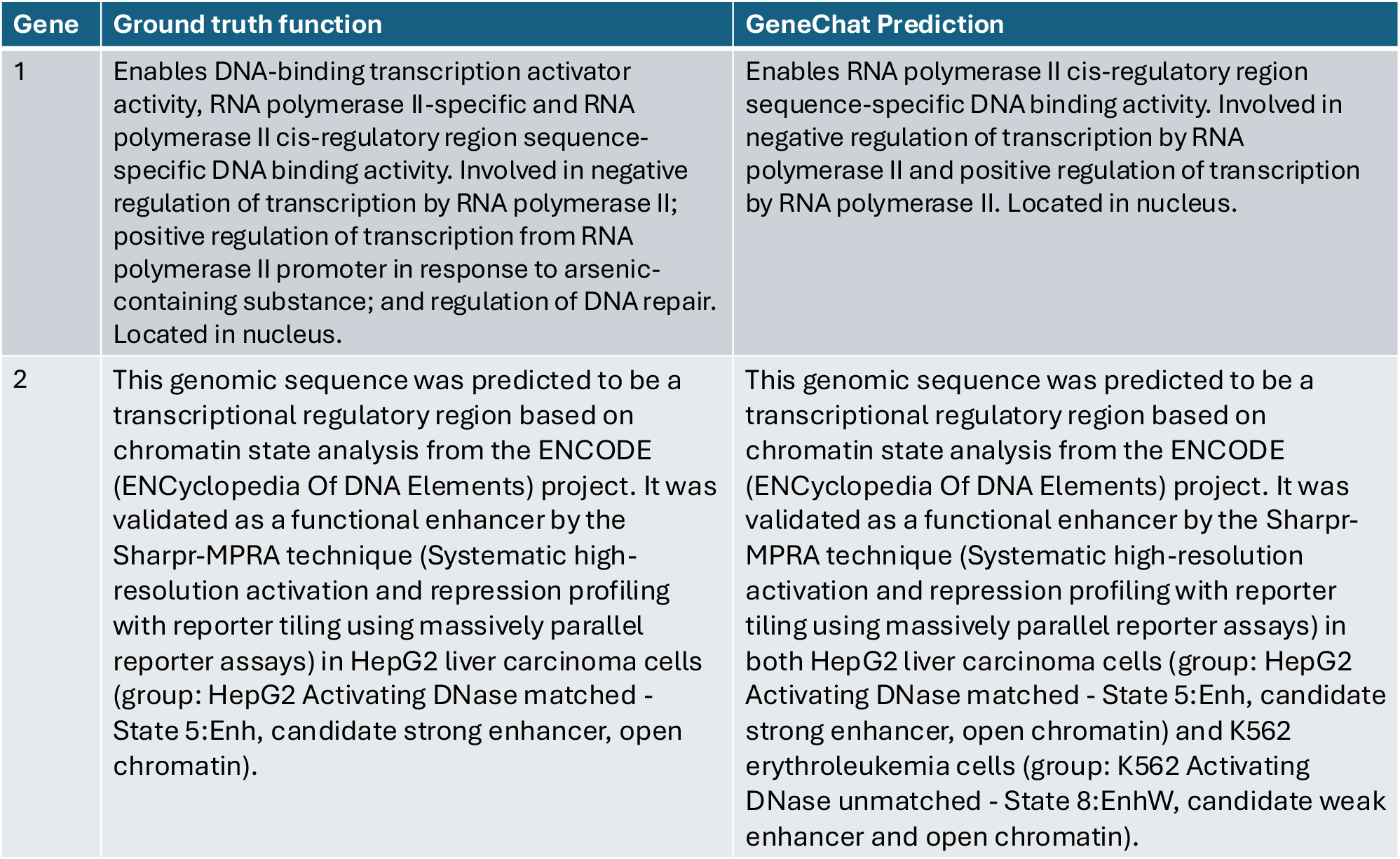
Examples of gene function predictions generated by GeneChat, compared against ground truth annotations.

## Discussion

GeneChat represents a significant advancement in computational genomics by introducing a free-form, language-based paradigm for gene function prediction. Unlike traditional models that frame function prediction as a multi-label classification task over predefined ontologies, GeneChat generates detailed, natural language descriptions that capture the complexity, context-dependence, and multi-functionality of genes. This flexibility enables more nuanced and interpretable outputs, bridging the gap between structured prediction and human-readable biological insight. The model’s architecture — integrating a genomic encoder with a large language model via an adaptor — demonstrates the value of combining domain-specific sequence representations with general-purpose linguistic reasoning. This design enables GeneChat to generalize across diverse gene types and organisms while remaining grounded in biologically meaningful features.

The superior performance of GeneChat over GPT-4o can be attributed to its domain-specific architecture and multi-modal design, which are tailored for gene function prediction. Unlike GPT-4o, which is a general-purpose large language model trained primarily on natural language text, GeneChat integrates a dedicated gene encoder, DNABERT-2, pre-trained on genomic sequences to capture biologically relevant features such as long-range dependencies and nucleotide-level patterns. This encoder enables the model to effectively interpret and represent the complex structure of DNA sequences—something GPT-4o is not designed to handle, as evidenced by its inability to generate meaningful outputs when provided with raw nucleotide sequences. Furthermore, GeneChat uses an adaptor module to align the high-dimensional gene embeddings with the latent space of a large language model, ensuring that sequence-derived representations are semantically compatible with natural language reasoning. This fusion allows GeneChat to leverage both biological context and linguistic knowledge when generating functional descriptions. In contrast, GPT-4o relies solely on textual cues, such as gene names, and lacks direct access to the sequence-level information essential for accurate and detailed function prediction. This limitation results in outputs that are often generic or inconsistent with the true biological role of the gene. The performance gap, reflected in substantially higher BLEU and METEOR scores for GeneChat, underscores the value of incorporating structured biological modalities and targeted pretraining when adapting large language models to specialized tasks in genomics.

The potential applications of GeneChat span multiple areas of biomedical research. In genomics, it can be used to annotate genes from newly sequenced organisms, particularly in cases where experimental data or close homologs are lacking. In functional genomics, GeneChat can assist in prioritizing candidate genes for experimental validation by providing interpretable hypotheses about their roles. Its ability to process raw nucleotide sequences also positions it as a useful tool in variant interpretation, where understanding the functional impact of non-coding or novel sequences is critical. In education and communication, the model’s natural language outputs offer a user-friendly interface for explaining gene functions to researchers, clinicians, and students. More broadly, GeneChat’s architecture serves as a blueprint for building multi-modal foundation models for other biomolecular tasks, such as enhancer annotation, non-coding RNA characterization, or protein–DNA interaction prediction, highlighting its potential as a general framework for biological sequence understanding.

Despite its strengths, GeneChat has several limitations that warrant consideration. First, the quality of its predictions is inherently constrained by the training data, which are derived from publicly available gene annotations that may be incomplete, inconsistent, or biased toward well-studied genes and organisms. As a result, the model may underperform when predicting functions for genes with limited or noisy training examples, particularly in non-model species. Second, while GeneChat generates free-form natural language outputs, these predictions are not guaranteed to be factually accurate or experimentally validated. The model may produce confident-sounding but incorrect statements, reflecting known challenges with hallucination in large language models. Third, although DNABERT-2 captures long-range dependencies in genomic sequences, the need to split long sequences into fixed-size chunks with overlapping windows may still limit the model’s ability to reason over very large regulatory or structural contexts. Fourth, GeneChat is currently limited to predicting gene function from sequence alone; it does not incorporate other valuable data modalities such as gene expression, chromatin accessibility, or protein–protein interactions, which are often essential for resolving functional ambiguity. Finally, the interpretability of the underlying reasoning process remains limited, as the model functions as a black box, making it difficult to trace specific predictions to particular sequence features or learned representations. Addressing these limitations will be crucial for enhancing the reliability, generalizability, and utility of free-form gene function predictors in real-world biological applications.

Future work on GeneChat will focus on enhancing its predictive accuracy, interpretability, and generalizability across diverse biological contexts. One key direction is the integration of additional data modalities beyond nucleotide sequences, such as gene expression profiles, epigenetic marks, protein–protein interaction networks, and evolutionary conservation scores. Incorporating such information could enable the model to make more context-aware predictions, particularly for genes with regulatory or condition-specific functions. Another area of development is improving model interpretability. Although GeneChat produces human-readable outputs, the internal reasoning behind its predictions remains opaque. Future extensions could incorporate attention visualization, saliency mapping, or structured explanation generation to increase transparency and support hypothesis generation. Scaling to larger genomic contexts is also a priority, particularly for predicting functions that depend on long-range regulatory interactions. Techniques such as hierarchical encoding or memory-augmented architectures may help address current sequence length limitations. Additionally, we aim to evaluate GeneChat in zero-shot and cross-species scenarios, testing its ability to generalize to previously unseen organisms or gene families. From a deployment perspective, adapting GeneChat into interactive tools — such as web-based platforms or APIs — could facilitate broader accessibility for researchers and educators. Finally, establishing benchmarks with experimental validation will be essential to assess the real-world utility of GeneChat predictions and to build trust in its applications in genomics, precision medicine, and biotechnology.

## Methods

### Data

The training data for GeneChat was curated from the National Center for Biotechnology Information (NCBI) database (15). It includes comprehensive genomic information spanning a wide range of organisms and contains 50,248 genes. Each gene is annotated with multiple attributes, including its nucleotide sequence, a detailed functional description, associated organism, gene type, chromosomal location, official symbol, full name, and exon count. The function description provides insight into the gene’s biological role, regulatory properties, and functional significance. The dataset was randomly partitioned into training and test sets using a 95:5 ratio. To train GeneChat, the nucleotide sequence of each gene was used as input, while the corresponding function description served as the target output. The training prompt used was: please predict the function of this gene.

### Model

Our model consists of three key components (Figure 1): a gene encoder, a large language model (LLM), and an adaptor that bridges the two. The gene encoder processes the input genomic sequence and generates a representation that captures long-range dependencies within the gene. The adaptor transforms this representation into a format compatible with the LLM’s latent space, enabling effective integration. Once aligned, the LLM incorporates both the genomic representation and the textual prompt to generate a free-form prediction of the gene’s function. For the gene encoder, we employed DNABERT-2 (12), a domain-specific pretrained model optimized for genomic sequences. As the language model component, we used Vicuna (13), a fine-tuned variant of LLaMA-2 (14), known for its strong instruction-following capabilities.

DNABERT-2 is an improved version of DNABERT (18), specifically developed for genomic sequence analysis using transformer-based architectures. While conventional transformers (9) are constrained by quadratic computational complexity and limited sequence lengths, DNABERT-2 incorporates architectural and algorithmic optimizations that enable modeling of longer DNA sequences—an essential capability for capturing long-range dependencies in genomic regions. A central innovation in DNABERT-2 is its dynamic k-mer strategy. Unlike the fixed k-mer tokenization used in the original DNABERT, DNABERT-2 learns variable-length k-mers, preserving single-nucleotide resolution while improving representational efficiency. This is particularly important for detecting single nucleotide polymorphisms and characterizing mutations that affect gene function. The model also introduces parameter-efficient transformer variants to reduce memory usage and computational cost, and incorporates genomic-position-aware embeddings and enhanced attention mechanisms for improved modeling of long-range sequence interactions. To further enhance training stability and performance, DNABERT-2 adopts a sequence length warm-up strategy that incrementally increases input length during training. This mitigates gradient instability and accelerates convergence, enabling effective learning from large-scale genomic datasets.

Vicuna is an open-source chatbot model fine-tuned from the LLaMA-2 (14) framework using approximately 70,000 user-shared conversations collected from ShareGPT.com (13). It retains the architecture of LLaMA-2-13B, consisting of 40 transformer layers, 40 attention heads, and a hidden embedding size of 5120. Vicuna is designed to generate coherent, contextually appropriate outputs, making it well-suited for applications such as conversational agents, text summarization, and open-ended content generation. The model is available in multiple variants, including 7B and 13B, which differ in model depth, width, and attention dimensionality to balance performance with computational efficiency. Instruction tuning further aligns the model with task-specific prompts, while the output layer computes token probabilities for generating fluent and relevant responses. These capabilities make Vicuna a strong backbone for language modeling tasks requiring instruction-following and free-form generation.

In GeneChat, given an input gene sequence *x*_*g*_, the DNABERT-2 encoder produces a contextual embedding *h*(*x*_*g*_) ∈ℝ^*l×*768^, where *l* is the length of the input sequence. Since *l* typically exceeds the context window size of DNABERT-2, we apply a pooling operation over non-overlapping windows of size *k*, reducing the sequence length to *l/k* and yielding a pooled embedding of shape ℝ ^(*l/k*)*×*256^. To project these pooled embeddings into the latent space of the LLM, we apply a linear adaptor layer *W* ∈ ℝ^256*×*5120^. The transformed embedding is given by:

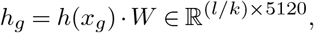

where *h*_*g*_ represents the gene representation aligned to the LLM’s latent embedding space. To integrate the gene embedding with the textual prompt, we format the input and output of the LLM following the conversational style used by Vicuna. Specifically, each training instance is structured as follows:

- (LLM Input) Human: <Gene> GeneHere </Gene>Prompt Assistant:
- (LLM Response) Answer

Here, the placeholders `Prompt` and `Answer` are replaced with the actual prompt and corresponding function description from the training triplet (*x*_*g*_, *x*_*prompt*_, *y*). The entire input of the LLM, excluding the token GeneHere, is referred to as the *auxiliary prompt*, which includes the literal tokens <, >, and /. We denote the tokenized auxiliary prompt as *x*_aux_, and use the LLM’s embedding layer to convert it into a sequence of embeddings *h*_aux_. To inject the gene-specific information, we replace the placeholder token `GeneHere` with the gene embedding *h*_*g*_, which is produced by applying the adaptor to the pooled output of the DNABERT-2 encoder. The final input to the LLM is a concatenation of *h*_aux_ and *h*_*g*_, forming a unified input sequence used to generate the textual prediction of the gene’s function.

The GeneChat model is trained using a causal language modeling objective, where the goal is to predict each token in the output sequence conditioned on the preceding context. Specifically, the model is optimized to maximize the log-likelihood of the target answer tokens. In our formulation, only the `Answer` portion of the LLM output is used to compute the loss. To enable the model to learn when to terminate generation, we append a special end-of-sequence token to each answer during training. Let **x**_*a*_ denote the answer sequence of length *l*, **x**_*g*_ the gene sequence input, and **x**_aux_ the tokenized auxiliary prompt. The conditional probability of generating **x**_*a*_ given **x**_*g*_ and **x**_aux_ is defined as:

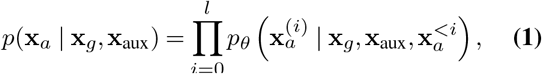

where 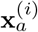 denotes the *i*-th token of the answer and 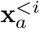 denotes all preceding tokens. The model parameters are denoted by *θ*. During training, the objective is to minimize the negative log-likelihood of this conditional probability.

### Training details of GeneChat

Since DNABERT-2 cannot process sequences as long as 160,000 nucleotides, we partition each input sequence into smaller chunks of 512 nucleotides. To preserve contextual continuity across segments, a 10-nucleotide overlap is maintained between consecutive chunks.

For the gene encoder (DNABERT-2), we applied full fine-tuning by updating all pretrained parameters during training. For the language model component, Vicuna-13B, we employed Low-Rank Adaptation (LoRA) (19), a parameter-efficient fine-tuning method that injects trainable low-rank matrices into the attention and feedforward layers while keeping the original weights frozen, thereby reducing memory and computational overhead. In our setup, LoRA was applied to the query and value projection matrices, with a rank of 8, a scaling factor (`alpha`) of 16, and a dropout rate of 0.05. The adaptor module, which aligns the gene representation with the LLM’s latent space, was trained from scratch.

Optimization was performed using the Adam optimizer (20) with *β*_1_ = 0.9, *β*_2_ = 0.999, and a weight decay of 0.05. A cosine learning rate decay schedule was used, with a peak learning rate of 1*×* 10^−4^, a linear warm-up over the first 2000 steps, and a minimum learning rate of 1 *×* 10^−6^. Due to memory constraints, we adopted a mini-batch size of 1 per GPU and accumulated gradients over 8 steps, resulting in an effective batch size of 8. The model was trained for a total of 170,000 steps.

### Evaluation metrics

We evaluated the quality of the generated gene function descriptions using BLEU (16) and METEOR (17) scores, two widely used metrics in natural language generation tasks. The BLEU (Bilingual Evaluation Understudy) score measures the overlap between predicted and reference texts using modified n-gram precision, along with a brevity penalty to penalize excessively short outputs. It is computed as:

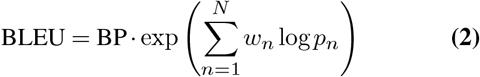

where *p*_*n*_ is the modified precision for n-grams of length *n*, and *w*_*n*_ is the corresponding weight (typically uniform). The brevity penalty (BP) discourages short predictions and is defined as:

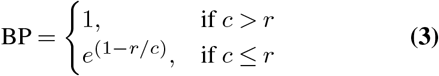

where *c* is the length of the predicted text and *r* is the length of the reference text.

The METEOR (Metric for Evaluation of Translation with Explicit ORdering) score complements BLEU by incorporating recall, synonym matching, stemming, and word order. It is designed to better align with human judgment. The score is computed using a harmonic mean of unigram precision (*P*) and recall (*R*), weighted by a fragmentation penalty that captures word order differences:

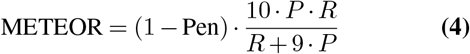

where Pen is a penalty based on the number of chunks — contiguous matched subsequences — reflecting word order dissimilarity. These metrics provide complementary views of output quality: BLEU emphasizes n-gram precision and length fidelity, while METEOR rewards recall, alignment, and semantic similarity.

### Related work

Recent advances in genomic foundation models have significantly advanced our ability to decode and interpret DNA sequences. The Nucleotide Transformer family (21) represents a major milestone, with models ranging from 50 million to 2.5 billion parameters pretrained on thousands of human and multispecies genomes. These models excel in zero-shot prediction of genetic variant effects and achieve state-of-the-art performance on tasks such as promoter identification and transcription factor (TF) binding prediction. By incorporating evolutionary context from diverse organisms, they capture conserved regulatory elements and enable accurate molecular phenotype prediction, even in low-data settings.

DNABERT (18) was one of the earliest bidirectional transformer models pretrained on human genomic sequences. It pioneered task-specific fine-tuning for predicting regulatory features such as splice sites and TF binding sites. Its attention maps also offer interpretable insights into nucleotide-level importance, facilitating motif discovery. DNABERT-2 (22) extended this line of work by introducing a multi-modal architecture that integrates DNA breathing dynamics — transient local strand separations — using a cross-attention mechanism. This enhancement improved cross-species TF binding predictions and enabled the identification of disease-associated non-coding variants, increasing the model’s biological relevance.

To address long-range dependencies in genomic sequences, HyenaDNA (23) replaced traditional attention mechanisms with Hyena operators, supporting context lengths of up to one million nucleotides at single-nucleotide resolution. Despite having fewer parameters, HyenaDNA outperformed attention-based models such as Nucleotide Transformer on tasks like chromatin accessibility profiling and species classification, while training up to 160× faster. This efficiency makes it well-suited for whole-genome analysis.

Together, these models illustrate the growing versatility of genomic foundation architectures. While Nucleotide Transformer and DNABERT variants emphasize interpretability and cross-species generalization, HyenaDNA prioritizes scalability for ultra-long genomic sequences. Their complementary strengths suggest promising directions for hybrid models in future genomic AI applications.

## Data availability

All data used in this study are available at https://drive.google.com/drive/folders/1g0Pe0HxfzdhXWbG54rkd-Iya7c6wYZdO?usp=sharing.

## Code availability

The source code of this work is available at https://github.com/Shashi-Sekar/GeneChat.

## References

1. Predrag Radivojac, Wyatt T Clark, Tal Ronnen Oron, Alexandra M Schnoes, Tobias Wittkop, Artem Sokolov, Kyle Graim, Christopher Funk, Karin Verspoor, Asa Ben-Hur, et al. A large-scale evaluation of computational protein function prediction. Nature methods, 10(3): 221–227, 2013.

2. Iddo Friedberg. Automated protein function prediction—the genomic challenge. Briefings in bioinformatics, 7(3):225–242, 2006.

3. Stephen F Altschul, Warren Gish, Webb Miller, Eugene W Myers, and David J Lipman. Basic local alignment search tool. Journal of molecular biology, 215(3):403–410, 1990.

4. Joshua M Stuart, Eran Segal, Daphne Koller, and Stuart K Kim. A gene-coexpression network for global discovery of conserved genetic modules. Science, 302(5643):249–255, 2003.

5. Christian von Mering, Roland Krause, Berend Snel, Marina Cornell, Stephen G Oliver, Stanley Fields, and Peer Bork. Comparative analysis of gene expression and phenotypes between knockout mice and human diseases. Nature, 417(6886):399–403, 2002.

6. Michael Ashburner, Catherine A Ball, Judith A Blake, David Botstein, Heather Butler, J Michael Cherry, Allan P Davis, Kara Dolinski, Suzanna S Dwight, Janan T Eppig, et al. Gene ontology: tool for the unification of biology. Nature genetics, 25(1):25–29, 2000.

7. Zhen Huo and Pengtao Xie. Proteinchat: A multi-modal large language model for protein function prediction. bioRxiv, 2024. doi:10.1101/2024.01.01.123456.

8. Tom B Brown, Benjamin Mann, Nick Ryder, Melanie Subbiah, Jared Kaplan, Prafulla Dhariwal, Arvind Neelakantan, Pranav Shyam, Girish Sastry, Amanda Askell, et al. Language models are few-shot learners. Advances in neural information processing systems, 33: 1877–1901, 2020.

9. Ashish Vaswani, Noam Shazeer, Niki Parmar, Jakob Uszkoreit, Llion Jones, Aidan N Gomez, Łukasz Kaiser, and Illia Polosukhin. Attention is all you need. Advances in neural information processing systems, 30, 2017.

10. Zheng Lin, Hayrettin Akin, Bo Ding, Maurice Sherman, Mallory Mak, Caleb Lareau, Yaping Liu, Ajay Sathe, Pirouz Mehdipour, Xiang Li, et al. A language model for dna sequences reveals connections to chromatin and transcription. Nature Biotechnology, 40(8):1202–1210, 2022.

11. Alexander Rives, Joshua Meier, Tom Sercu, Siddhartha Goyal, Zeming Lin, Jason Liu, Dmytro Guo, Myle Ott, C Lawrence Zitnick, Jerry Ma, and Rob Fergus. Biological structure and function emerge from scaling unsupervised learning to 250 million protein sequences. Proceedings of the National Academy of Sciences, 118(15), 2021.

12. Zhihao Zhang, Peng Li, Chong Yang, Xingyu Wang, Jianzhu Li, and Zhiyong Lu. Dnabert-2: Effective dna language pre-training with longer k-mers. bioRxiv, 2023. doi: 10.1101/2023.04.14.536902. Available at https://www.biorxiv.org/content/10.1101/2023.04.14.536902v1.

13. Wei-Lin Chiang, Zhuohan Xu, Hao Zhao, Shang-Wen Zhuang, Xinyang Lin, Lianmin Deng, Ying v. d. Gablentz, et al. Vicuna: An open-source chatbot impressing gpt-4 with 90%* chatgpt quality. arXiv preprint arXiv:2306.05685, 2023.

14. Hugo Touvron, Louis Martin, Kevin Stone, Peter Albert Babaei, Thibaut Lavril, Daniel Zwemer, et al. Llama 2: Open foundation and fine-tuned chat models. Meta AI Technical Report, 2023.

15. Eric W Sayers, Mark Cavanaugh, Karen Clark, James Ostell, Kim D Pruitt, and Ilene Karsch-Mizrachi. Database resources of the national center for biotechnology information. Nucleic Acids Research, 49(D1):D10–D17, 2021. doi: 10.1093/nar/gkaa892.

16. Kishore Papineni, Salim Roukos, Todd Ward, and Wei-Jing Zhu. Bleu: a method for automatic evaluation of machine translation. In Proceedings of the 40th annual meeting of the Association for Computational Linguistics, pages 311–318, 2002.

17. Satanjeev Banerjee and Alon Lavie. Meteor: An automatic metric for mt evaluation with improved correlation with human judgments. In Proceedings of the ACL workshop on intrinsic and extrinsic evaluation measures for machine translation and/or summarization, pages 65–72, 2005.

18. Xin Huang Jie Jiang Yonghong Tian Jianguo Zhang Jiayu Ji, Zirui Wang. Dnabert: Pretrained bidirectional encoder representations from transformers for dna-language. Bioinformatics, 37(15):2112–2120, 2021. doi: 10.1093/bioinformatics/btab089.

19. Edward J. Hu, Yelong Shen, Phillip Wallis, Zeyuan Allen-Zhu, Yuanzhi Li, Shean Wang, and Weizhu Chen. Lora: Low-rank adaptation of large language models, 2021.

20. Diederik P. Kingma and Jimmy Ba. Adam: A method for stochastic optimization, 2017.

21. Aurélien Bonnafoux, Gianluca Bontempi, et al. The nucleotide transformer: Building and evaluating robust foundation models for human genomics. bioRxiv, 2023. doi: 10.1101/2023.01.11.523679. Preprint. Version 1. Available at: https://www.biorxiv.org/content/10.1101/2023.01.11.523679v1.

22. Zhihan Zhou, Yanrong Ji, Weijian Li, Pratik Dutta, Ramana Davuluri, and Han Liu. Dnabert-2: Efficient foundation model and benchmark for multi-species genome. 2024.

23. Marjan Faizi et al. Eric Nguyen, Michael Poli. Hyenadna: Long-range genomic sequence modeling at single nucleotide resolution. 2024.

